# GPhase: Greedy Approach for Accurate Haplotype Inferencing

**DOI:** 10.1101/073379

**Authors:** Kshitij Tayal, Naveen Sivadasan, Rajgopal Srinivasan

**Affiliations:** Life Sciences Division, TCS Innovation Labs, Hyderabad 500081, India

## Abstract

We consider the computational problem of phasing an individual genotype sample given a collection of known haplotypes in the population. We give a fast and accurate algorithm GPhase for reconstructing haplotype pair consistent with input genotype. It uses the coalescent based mutation model of Stephens and Donnelly (2000). Computing optimal solution under this model is expensive and our algorithm uses a greedy approximation for fast and accurate estimation. Our algorithm is simple, efficient and has linear time and space complexity. Experiments on real datasets revealed improved gene level phasing accuracy for GPhase tool compared to other widely used tools such as SHAPEIT, Beagle, MaCH and Impute2. On simulated data, GPhase tool was able to phase samples each containing more than 1700 markers with high accuracy. GPhase can be used for gene level phasing of individual samples using publicly available haplotype datasets such as HapMap data or 1000 genome data. This finds applications in studies on recessive Mendelian disorders where parent data is lacking. GPhase is freely available for download and use from https://github.com/kshitijtayal/GPhase/.

## I. Introduction

Identification of genomic variants is known as variant calling. Since functional consequences very often depend on having two or more such variations either as part of the same or different haplotypes, it is critical to establish the relationship between SNPs as to whether they occur as part of the same haplotype or different haplotypes. Haplotype reconstruction from genotype data is known as phasing. In this work we consider the problem of phasing an individual genotype given a collection of known haplotypes.

Diploid organisms such as humans carry two homologous copies of each chromosome, one from each parent. Mutations and recombinations results in haplotype diversity in a population. In this work, we consider only biallelic genotypes with SNPs. Biallelic haplotypes can be represented as binary vectors, where a 0 and a 1 at a given SNP location (locus) indicates reference and alternative alleles respectively. Haplotypes in Fig. 1 can thus be represented by (1, 0, 0, 0, 0, 1) and (1, 1, 0, 1, 1, 0) and the genotype is represented by (1/1, 0/1, 0/0, 0/1, 0/1, 0/1). First and third locus in this case are homozygous and rest are heterozygous. Since there are 4 heterozygous sites in this genotype, there are in total 2^4^ haplotypes consistent with it. Thus, the solution space of haplotypes consistent with a given genotype has exponential dependence on the number of heterozygous sites.

**Fig. 1.**
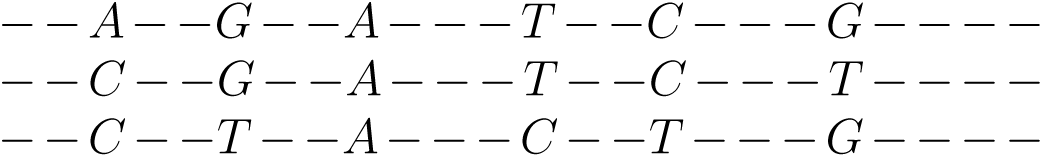
First sequence is reference sequence. Other two sequences are two copies of a chromosome. Non SNP positions are marked by - while the rest are SNPs. There are 6 SNPs in total.

The whole gamut of haplotype determination algorithms broadly falls under either haplotype assembly or haplotype inference. Haplotype assembly uses sequence reads directly to determine constituent haplotypes whereas haplotype inference use genotype data to infer haplotypes. Our work falls under the latter.

Clark’s algorithm (1990) [3] was one of the first computational techniques used for haplotype inferencing. It follows parsimony approach where it attempts to restrict the number of distinct haplotypes observed in the sample. It performs reasonably well on instances with small set of markers, however its performance suffers if markers are sparsely connected. Expectation-Maximization (EM) based algorithms (1995) [6], [8], [12] assigns alleles to haplotypes with high likelihood using estimated values for population haplotype frequencies, typically assuming uniform prior for haplotypes. EM approaches are computationally expensive in case of large number of heterozygous sites.

Approximate coalescent based methods, where new haplotypes are viewed as being derived from existing ones through mutation and recombination, achieved significant improvements in phasing quality. PHASE(v1.1) (2001) [18] was the first statistical method based on approximate coalescent based mutation model of Stephens and Donnelly [16] which identifies new haplotypes as derivative of old haplotypes with mutations. It uses Gibbs sampling to sample from the underlying Markov chain on the haplotype solution space. It exhibited superior accuracy than Clarks and EM but suffered from very high computational overhead. PHASE(v2.1.1) (2005) [17] used coalescent with recombination model to account for linkage disequilibrium and recombination. Similar to PHASE(v1.1) it is also computationally expensive. PHASE [18], [17] was considered gold standard for accuracy [2] but lost its importance due to slow speed. fastPHASE (2006)[15] made it possible to phase thousands of heterozygous sites at a considerable speed at the cost of reduced accuracy than its predecessors. Speed up in fastPHASE is achieved by locally clustering the haplotype using Hidden Markov model (HMM).

Beagle (2007) [1], similar to fastPHASE, use HMM to infer haplotypes. Similar to fastPHASE, it performs local clustering of haplotypes at each position, but allowing different number of clusters at each marker. Haplotype clusters form HMM states here. Impute2 (2009) [9] and MaCH (2010) [11] can operate on large data sets than what PHASE can deal with and simultaneously achieving greater accuracy than fastPHASE. They can be used for imputation of untyped variants as well. SHAPEIT (2012)[5] is an enhancement over Impute2 and MACH that scales linearly in the number of SNPs. It is lot more faster, accurate and uses less memory as compared to other two. Availability of large amounts of public haplotype data such as 1000 genome data [4] and HapMap data [7] have contributed to further improvements in phasing accuracy. We refer the reader to [2] for a review on various phasing algorithms.

### A. Our Contribution

We consider the problem of phasing an individual genotype sample given a collection of known haplotypes in the population. The known haplotype collection could for instance be the publicly available haplotype data such as HapMap data [7] or 1000 genome data[4]. We give a fast and accurate algorithm GPhase for reconstructing a haplotype pair consistent with input genotype. Our algorithm is based on the coalescent based mutation model of Stephens and Donnelly [16] (without recombination) which is used in PHASE (v1.1) [18]. Computing optimal solution under this model is expensive and our algorithm uses greedy approximation for fast and accurate estimation. Our algorithm is simple, efficient and has linear time and space complexity. It constructs the solution incrementally while maintaining a collection of top *k* candidate solutions at each step and finally uses the top most solution as its solution. Experiments on real datasets revealed improved gene level phasing accuracy for GPhase tool compared to SHAPEIT, Beagle, MaCH and Impute2. On simulated data, GPhase tool was able phase samples each containing more than 1700 markers with high accuracy. Our algorithm can be viewed as an instance based learner that, for each input sample, infers most probable solution from whole training data (known haplotype collection) using the mutation model. This is in contrast to other approaches that either use simpler mutation models or build simple generalized models such as HMM from the training data for faster inferencing, which results in information/accuracy loss. GPhase tool can be used for gene level phasing of individual samples using publicly available haplotype datasets such as 1000 genome data. This has applications in studies on recessive Mendelian disorders in the absence of parent data. The tool is freely available for download and use from https://github.com/kshitijtayal/GPhase/.

## II. Algorithm

Notations: We use 0 and 1 to represent two alleles of a biallelic locus. A haplotype *h* is specified over *l* loci by an *l*-dimensional binary vector. A genotype *G* characterized over *l* loci is also an *l*-dimensional vector from {0, 1, 2}^*l*^, where 0, 1 and 2 are counts of the allele coded as 1. It is assumed that genotype data has no missing information. Let *h*(*i*) and *G*(*i*) denote the *i*th locus of haplotype *h* and genotype *G* respectively. Let *h*(*i* : *j*) and *G*(*i* : *j*) for *i* ≤ *j* denote the sequence of loci *i, i* + 1, …, *j* in *h* and *G* respectively. Pair of haplotypes (*h*_1_, *h*_2_), called diploid is said to be consistent with genotype *G* if *G*(*i*) = *h*_1_(*i*) + *h*_2_(*i*) for all *i* ∈ {1, …, *l*}. Let *A_n_* = {*a*_1_, …, *a_n_*} denote a multi-set of *n* haplotypes each consisting of *l* loci. We also refer to *A_n_* as an *n* × *l* haplotype matrix with rows numbered 1, …, *n* and columns numbered 1, …, *l* and *A_n_*(*i, j*) denoting the *j*th locus of haplotype *a_i_*. We may conveniently refer to *A_n_* as a set or as a matrix.

### A. Review of Mutation Model [16]

In this section we review relevant parts of the coalescent based model proposed by Stephens and Donnelly (2000) [16] which is used by PHASE (v1.1) [18] for haplotype inferencing. Our algorithm uses this model for inferencing. We refer the reader to [16], [18] for details. Let *A_n_* denote a multi-set of *n* known haplotypes each consisting of *l* loci. Let *π*(*h|A_n_*) denote the conditional distribution of observing an *l* loci haplotype *h* given *A_n_*. Stephens and Donnelly [16], proposes an approximation to *π*(*h|A_n_*) as

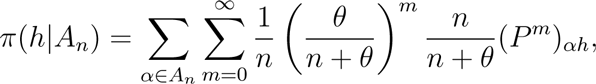

where *P* is the mutation matrix and *θ* is a scaled mutation rate. That is, *π*(*h|A_n_*) corresponds to the probability of choosing one haplotype say *α* uniformly at random from *A_n_* and applying *m* mutations to *α* to obtain *h*, where *m* is geometrically distributed with parameter *θ*/(*n* + *θ*). Since computing the above expression can be expensive, Stephens and Donnelly [16] makes further simplifying assumption that each locus mutates independently at rate *θ*/2 and thus with a total rate of *lθ*/2 and according to a 2 × 2 transition matrix (for biallelic case) *P*. In this simplified model, a haplotype *α* is chosen uniformly at random from *A_n_* and thereafter *m* locations (with repetition) are chosen uniformly from *α* and mutated according to mutation matrix *P*, where *m* is geometrically distributed with parameter *lθ*/(*n* + *lθ*). Using properties of Poisson distribution, this process can be equivalently viewed as drawing a time *t* from exponential distribution with rate parameter 1 and applying *m_i_* mutations to each locus *i* ∈ {1, …, *l*} where *m_i_* values are independent and are Poisson distributed with parameter *θt/n*. Mutations at each locus are again as per the mutation matrix *P*. This yields the following modified expression

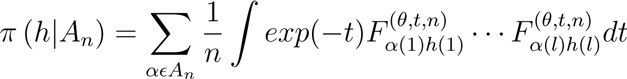

where

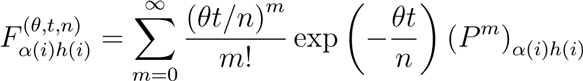

The integral in the above expression is further approximated using Gaussian quadrature to finally obtain

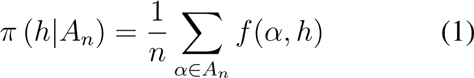

where

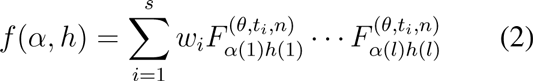

Here *t*_1_, …, *t_s_* are the quadrature points and *w*_1_, …, *w_s_* are the quadrature weights (marginal probabilities). Matrices 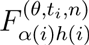 can be precomputed where the infinite sum can be well approximated by a finite sum of a large number of terms.

For a haplotype pair *H* = (*h*_1_, *h*_2_) we have

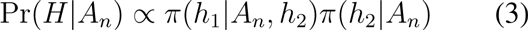

In fact, for large *n*, finding *H* = (*h*_1_, *h*_2_) that maximizes eq. (3) can be approximated by finding *H* that maximizes

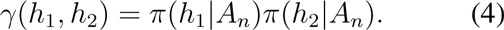

PHASE algorithm uses the above model to phase a collection of genotypes *G* = {*G*_1_, …, *G_n_*} [18]. Their algorithm uses Gibbs sampling where in each iteration, a random genotype *G_i_* from *G* is phased assuming all remaining genotypes in *G* are correctly phased. If *H*_−*i*_ denote the reconstructed haplotypes for all genotypes in *G* except *G_i_*, then most probable haplotype *h_i_* consistent with *G_i_* is inferred using equation (1) for *π*(*h|H*_−*i*_) and is selected as a haplotype for *G_i_*. The process is repeated until the underlying Markov chain on the solution space mixes.

Size of the set of haplotypes {*h_i_*} consistent with a genotype *G_i_* has exponential dependency on the number of heterozygous loci in *G_i_*. Hence, |{*h_i_*}| can be very large in general and this makes finding most probable *h_i_* consistent with genotype *G_i_* computationally expensive. This issue is partially addressed in PHASE [18] by considering only few random heterozygous loci in *G_i_* in each iteration of Gibbs sampling and thereby restricting the size of {*h_i_*} for each updation [18]. However this can result in increased mixing time. Simplified alternative models were also considered in other tools such as SHAPEIT and fastPHASE for performance improvements. In the next section, we describe our algorithm that approximately solves the problem of computing a haplotype pair (*h*_1_, *h*_2_) consistent with genotype *G* that maximizes eq (4) given a collection of known haplotypes *A_n_*. Our algorithm runs in time linear in the number of heterozygous loci in *G*. We consider only biallelic genotypes.

### B. GPhase Algorithm

We describe GPhase algorithm to solve the following problem: given a collection *A_n_* of known haplotypes and an unphased genotype *G*, reconstruct the haplotype pair *H* = (*h*_1_, *h*_2_) that is consistent with *G* and maximizes *γ*(*h*_1_, *h*_2_) given by equation (4). If there is a small collection of genotypes to be phased given the known haplotype collection *A_n_*, then the algorithm can be used to phase each genotype in the collection separately. We describe a greedy algorithm to solve this problem approximately. Our experiments for gene level phasing show that the phasing quality achieved by our greedy approximation exhibit superior quality compared to other widely used phasing tools.

Let *l* denote the number of loci in *G* and in haplotypes belonging to *A_n_*. For brevity of notation, given *θ* and *n*, let

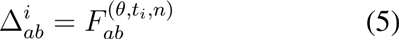

where *a, b* ∈ {0, 1}. We remark that 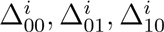 and 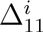 for *i* ∈ {1, …, *s*} are fixed for a given problem as *θ* and *n* are fixed. Given two haplotypes *α* and *β*, *f*(*α, β*) can be written as

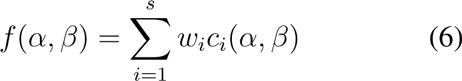

where

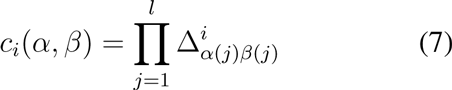

For *i* ∈ {1, …, *s*}, *c_i_*(*α, β*) can be interpreted as a ‘product similarity’ between *α* and *β* where for each locus *j*, depending on (*α*(*j*)*β*(*j*)) ∈ {00, 01, 10, 11}, a multiplicative term 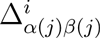 is contributed to *c_i_*(*α, β*). Function *f*(*α, β*) can hence be interpreted as the weighted sum of *s* similarity values *c*_1_(*α, β*), …, *c_s_*(*α, β*). It is easy to see that each locus contributes independently to similarity values *c_i_*(*α, β*) and the final weighted sum *f*(*α, β*) is invariant to the relative ordering of loci. It is also easy to verify that the same holds true for *π*(*h|A_n_*) and that the optimal solution that maximizes *γ*(*h*_1_, *h*_2_) is thus invariant any fixed permutation of columns of *A_n_* and loci of *G* where the same permutation is applied to both.

For ease of exposition, before describing the GPhase algorithm, we consider a related problem: given a collection *A_n_* of known haplotypes and an unphased genotype *G*, find haplotype *h* that is consistent with *G* and maximizes *π*(*h|A_n_*). We give a fast greedy algorithm GPhaseSingle (Algorithm 1) to find an approximate solution to this problem. We remark that this is a fundamental and computationally expensive sub-problem in some of the existing phasing tools. Our approach could hence be of use in other phasing methods as well. We will later extend this algorithm to solve the original problem. The algorithm considers one locus at a time and builds candidate solutions incrementally in a greedy fashion. It maintains its top *k* candidate solutions (for a fixed parameter *k*) that are updated after each incremental extension. The order in which loci of *G* is considered for phasing is governed by the column ordering of matrix *A_n_* resulting from the following *skewness permutation* of its columns.

#### Skewness permutation of A_n_, G

Let *A_n_*(*j*) be the *j* th column of matrix *A_n_*. Let (*j*) denote the absolute value of difference between total number of 1s and total number of 0s in column *A_n_*(*j*). Skewness permutation of the columns of matrix *A_n_* and correspondingly genotype *G* are done in the following manner. Let *L* ⊆ {1, …, *l*} denote the subset of heterozygous loci in *G* and let *l*′ = |*L*|. Reorder columns of *A_n_* such that its first *l* − *l*′ columns correspond to columns in {1, …, *l*} − *L* (corresponding to homozygous loci in *G*) in an arbitrary order. Remaining *l*′ columns correspond to columns in *L* arranged in the decreasing order of their corresponding *δ*(*j*) values. Hence, in the permuted matrix of *A_n_*, columns corresponding to homozygous loci in *G* appear first followed by columns corresponding to heterozygous loci in *G* in the decreasing order of their 0/1 skewness (*δ*(*j*) values).

By considering columns of the permuted matrix from left to right, loci (columns) with larger 0/1 skewness are considered earlier because phasing them can be done with lesser uncertainty. From now we assume that columns of *A_n_* (and correspondingly loci in *G*) are reordered based on its skewness permutation. Recalling the notations, let *h*(1 : *j*) for *j* < *l* denote a partial haplotype solution for loci 1, …, *j*. Let *π* (*h*(1 : *j*)|*A_n_*) denote the value of *π*(*h|A_n_*) when both *h* and *A_n_* are restricted to columns 1, …, *j*. The greedy algorithm maintains top-k partial solutions *h*_1_(1 : *j*), …, *h_k_*(1 : *j*), for a fixed parameter *k*, using a *k* element min heap data structure based on their *π* (*h_i_*(1 : *j*)|*A_n_*) values. Recalling that the first *l* − *l*′ loci of *G* are all the homozygous loci of *G* (after permutation), the heap is initialized with a single partial solution *h*(1 : *l*−*l*′) where *h*(*j*) = *G*(*j*) for *j* ∈ {1, …, *l* − *l*′}, with its corresponding value *π*(*h_i_*(1 : *l* − *l*′)|*A_n_*). Each candidate top-k partial solution *h*(1:*j*) from the heap is extended by one locus *j* + 1 in the following manner. Let *h*(1 : *j*)0 denote the sequence of length *j* + 1 obtained by appending 0 as the rightmost element to the sequence *h*(1 : *j*). Consider the two possible extensions *h*_0_ = *h*(1 : *j*)0 and *h*_1_ = *h*(1 : *j*)1 which are *j* + 1 length haplotypes obtained by appending 0 and 1 respectively to *h*(1 : *j*). Both *h*_0_ and *h*_1_ are considered for insertion into the top *k* heap of extended solutions based on their values *π*(*h*_0_(1 : *j* +1)|*A_n_*) and *π*(*h*_1_(0 : *j* + 1)|*A_n_*). Finally, after scanning all *l* loci, haplotype *h* with maximum value in the *k*-heap is output as the solution. Top *k* partial solutions are maintained instead of only the current best in order to handle situations where final top solutions are suboptimal for the partial solutions considered during extension steps. Larger values of *k* would improve the quality of final solution. These steps are given below as Algorithm 1. Parameters for the algorithm are mutation rate, quadrature points {*t*_1_, …, *t_s_*}, quadrature weights {*w*_1_, …, *w_s_*} and heap size *k*.

##### Algorithm 1. GPhaseSingle

~~~
   **Input:** *A_n_* and *G* on *l* loci.
   **Output:** Haplotype *h* consistent with *G*.
    /* Let *l*′ be the number of heterozygous loci in *G*. */
    Do skewness permutation of *A_n_, G*
    Initialize partial solution *h*(1 : *l* − *l*′) with
    *h*(*j*) = *G*(*j*) for *j* ∈ {1, …, *l* − *l*′}.
    Insert *h*(1 : *l* − *l*′) in 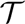 with value *π*(*h*(1 : *l* − *l*′)|*A_n_*).
    **for** *j* =*l* − *l*′ to *l* − 1 **do**
       *S* := Elements of 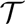.
       Empty 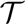.
       **for all** *h*(1 : *j*) in *S* **do**
          Consider extensions *h*_0_ = *h*(1 : *j*)0 and *h*_1_ = *h*(1 : *j*)1.
          Insert *h*_0_ in 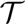 with value *π*(*h*_0_(1 : *j* + 1)|*A_n_*).
          Insert *h*_1_ in 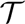 with value *π*(*h*_1_(1 : *j* + 1)|*A_n_*).
       **end for**
   **end for**
   **return** 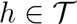 with maximum value.
 **end**
~~~

#### Implementation and Run time

Permutation of *A_n_* and *G* involves computing *δ*(*j*) values and sorting them, which can be achieved in *O*(*nl* + *l* log(*l*)) = *O*(*nl*) time. We note that since min heap 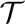 maintains top-*k* largest solutions, inserting a new element *h* in 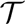 succeeds only if either heap size is less than *k* or if heap minimum is less than value of *h*, in which case the minimum element is replaced with *h*. In order to compute *π* (*h*(1 : *j*)|*A_n_*) efficiently, we maintain *k* two dimensional arrays of the form *B*[1..*n*][1..*s*], where *s* is the number of quadrature points, one such array for each of the top *k* partial solutions, with total *O*(*nk*) space. For each top *k* partial solution *h*(1 : *j*),*B* [*i*][*r*] entries in its corresponding *B* array store *c_r_*(*a_i_*(1 : *j*), *h*(1 : *j*)) values as given by eq. (7). That is, for a top *k* partial solution *h*(1 : *j*), we store its *s* component similarity values *c*_1_(), …, *c_s_*() between each haplo-type *a_i_* ∈ *A_n_* and *h*, both restricted to loci 1, …, *j*. From eq.(1) and eq.(6), it is easy to see that value of partial solution *π*(*h|A_n_*) is obtained from its corresponding *B* array as 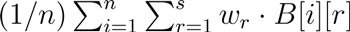. From eq. (7), it is straightforward to verify that if a partial solution *h*(1 : *j*) is extended by say 0, values in its *B*[*i*][*r*] array entries can be easily updated as either 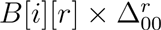 or 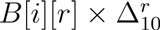 depending on whether *j* + 1st locus of *a_i_* is either 0 or 1. From Algorithm 1, it follows that the total cost of one iteration of the outer loop is *O*(*nk*) and the total run time is thus *O* (*nlk*). Thus, for fixed *k*, the algorithm runs in linear time and uses linear space.

#### GPhase algorithm

The final algorithm is obtained by straightforward modification of the previous GPhaseSingle algorithm. Instead of maintaining top *k* partial solutions which are single haplotypes, we maintain *k* haplotype pairs (*h*_1_(1 : *j*), *h*_2_(1 : *j*)) consistent with *G*(1 : *j*) as partial solutions. That is, *G*(*i*) =*h*_1_(*i*) + *h*_2_(*i*) for *i* = 1, …, *j*. Value of the partial solution is given by γ(*h*_1_(1 : *j*), *h*_2_(1 : *j*)) = *π*(*h*_1_(1 : *j*)|*A_n_*) · *π*(*h*_2_(1 : *j|A_n_*) (see eq.(4)). Extending a partial solution pair (*h*_1_(1 : *j*), *h*_2_(1 : *j*)) give rise to two new solution pairs viz. (*h*_1_(1 : *j*)0, *h*_2_(1 : *j*)1) and (*h*_1_(1 : *j*)1, *h*_2_(1 : *j*)0). Finally, the solution pair with maximum value is output. The final algorithm is given in Algorithm 2. Parameters for the algorithm remain the same.

By arguments similar to that for GPhaseSingle, it follows that GPhase require total *O*(*nlk*) time and uses *O*(*nk*) space. Hence it runs in linear time and uses linear space for fixed *k*.

##### Algorithm 2 GPhase

~~~
**Input:** *A_n_* and *G*.
**Output:** Haplotype pair (*h*_1_, *h*_2_) consistent with *G*.
 1: /* Let *l*′ be the number of heterozygous loci in *G*. */
 2: Do skewness permutation of *A_n_, G*
 3: Initialize partial solution (*h*_1_(1 : *l* − *l*′), *h*_2_(1 : *l* − *l*′)) with *h*_1_(*j*) = *h*_2_(*j*) = *G*(*j*) for *j* ∈ {1, …, *l* − *l*′}.
 4: Insert (*h*_1_(1 : *l* − *l*′), *h*_2_(1 : *l* − *l*′)) in 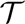 with value *γ*(*h*_1_(1 : *l* − *l*′), *h*_2_(1 : *l* − *l*′)).
 5: **for** *j* = *l* − *l*′ to *l* − 1 **do**
 6:    *S* := Elements of 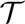.
 7:    Empty 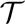.
 8:    **for all** (*h*_1_(1 : *j*), *h*_2_(1 : *j*)) in *S* **do**
 9:         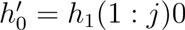 and 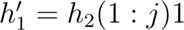
 10:        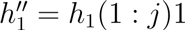 and 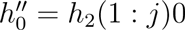
 11:       Insert 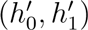 in 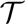 with value 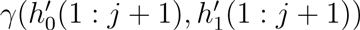.
 12:       Insert 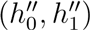 in 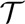 with value 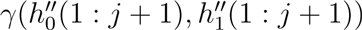.
 13:     **end for**
 14: **end for**
 15: **return** 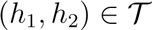 with maximum value.
**end**
~~~

## III. Results

We conducted experiments where we compared GPhase phasing quality with other existing tools. We performed experiments on both real datasets and simulated datasets. In particular, we compared GPhase output with phasing output of Beagle [1], MaCH [11], Impute2 [9] and SHAPEIT [5]. For comparison, we used the standard metrics of accuracy and switch accuracy [13]. Accuracy is defined as (*l* − 1 − *ms*)/(*l* − 1), where *l* is the number of heterozygous loci in the input genotype and *ms* is the number of mismatching loci between inferred haplotype and true haplotype. Switch accuracy is defined as (*l* − 1 − *sw*)/(*l* − 1) where *sw* is the number of switches to recover true haplotype from inferred haplotype.

In all our experiments, model parameters used by GPhase, namely mutation rate *θ*, quadrature points, quadrature weights and matrix *P* were same as that in PHASE implementation [17]. Value *θ* was set to 1/log(2*n*) and number of quadrature points were 2. The 2 × 2 transition matrix *P* required for pre-computing 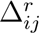 has entries *P*_00_ = *P*_11_ = 0 and *P*_01_ = *P*_10_ = 1. Value of *k* was set to 20.

### A. Real Dataset

For real dataset, we used in-house data that consisted of genotype samples from 10 unrelated individuals. For each individual, there were 1529 SNP markers from across chromosomes. These markers spanned 81 genes. There were 2463 heterozygous loci in total across all individuals. Constituent haplotypes for these individual genotype samples were already known from their nuclear family data. This was used for validation. We conducted separate phasing experiment on each gene level sample for each individual using the collection of known haplotypes extracted from 1000 genome data. Fig 2 shows the distribution of genes based on the average number of heterozygous loci in their corresponding genotypes (from 10 individuals).

**Fig. 2.**
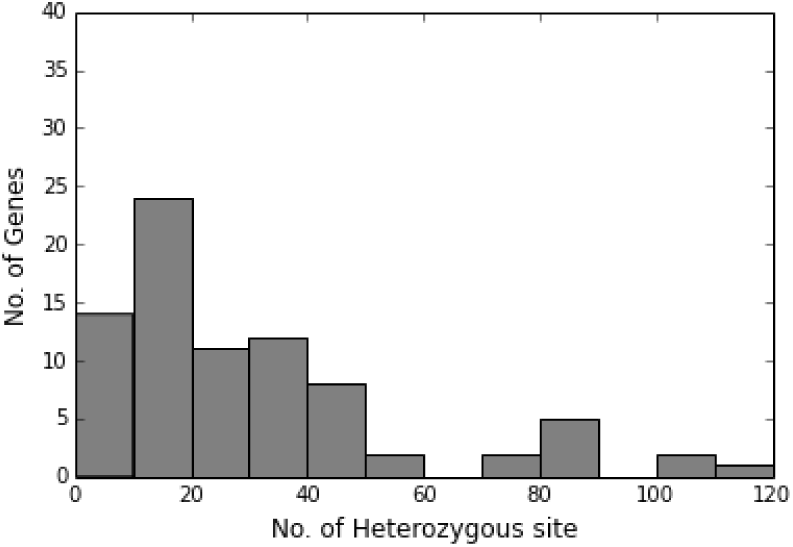
Distribution of gene counts based on the average no. of heterozygous sites for 10 individuals. Heterozygous count range is split into 12 equal sized intervals and no. of genes falling in each interval is plotted separately.

We used 1000 genome public dataset[4] to create the known haplotype set *A_n_*. In our experiments, *A_n_* consisted of 5008 individual haplotypes from 26 populations. All tools were run with default settings. We ran SHAPEIT on the inputs both under default recombination setting and under variable recombination setting. In the latter case, recombination data was provided using standard chromosome map files for human chromosomes [10]. Flag -no-mcmc was used with SHAPEIT and SHAPEIT RECOM.

Figures 4 and 3 provide accuracy plots for phasing outputs from all the candidate tools. They provide cumulative distribution of the percentage of genes (out of 81) whose mean accuracy and mean switch accuracy (across 10 individuals) are at least *w* for different accuracy values *w*. From these plots we see that GPhase phasing quality is superior to other tools for most of the plotted accuracy values. This is true in particular for higher accuracy values.

**Fig. 3.**
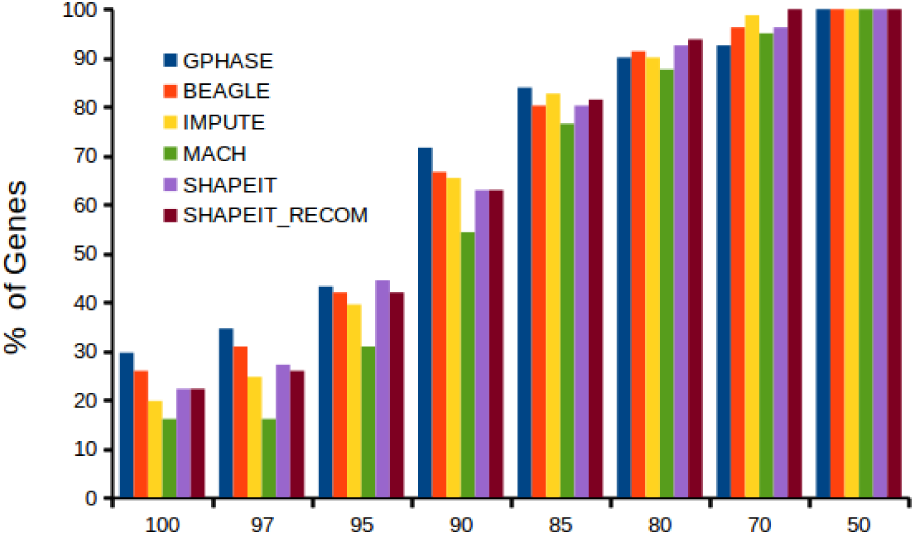
Tool-wise cumulative distributions where for different accuracy values, the percentage of genes whose mean switch accuracy is at least the specified accuracy value is plotted.

**Fig. 4.**
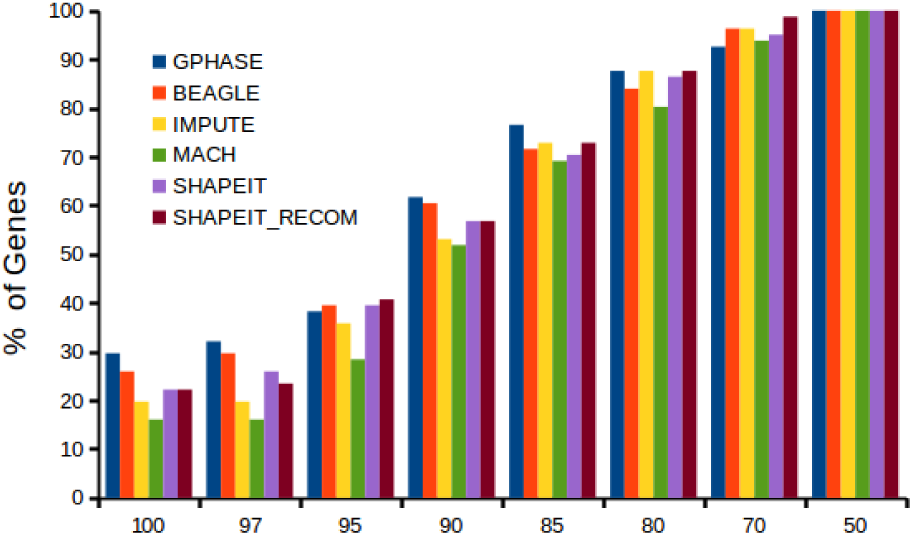
Tool-wise cumulative distributions where for different accuracy values, the percentage of genes whose mean accuracy is at least the specified accuracy value is plotted.

Figures 5 and 6 give plots of mean accuracy and mean switch accuracy of GPhase separately for inputs whose heterozygous loci count falls in different ranges. As evident from these plots, GPhase performs comparably well on inputs with wide range of heterozygous loci counts.

**Fig. 5.**
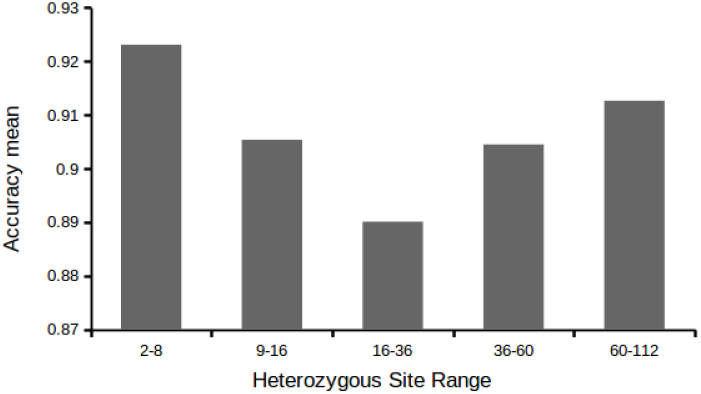
Mean accuracy of GPhase for genotype inputs falling under different heterozygous loci count ranges.

**Fig. 6.**
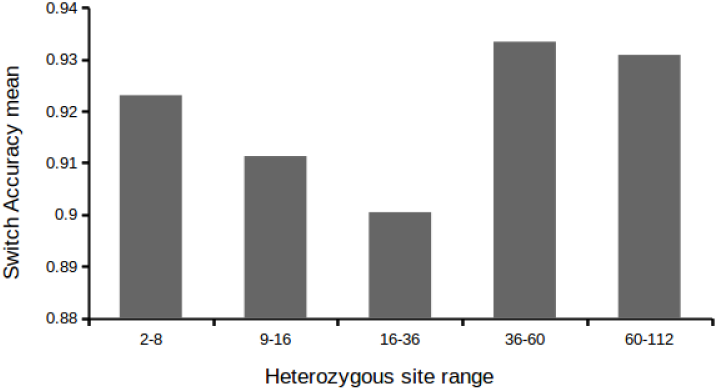
Mean switch accuracy of GPhase for genotype inputs falling under different heterozygous loci count ranges.

### B. Simulated Data

For simulated data generation, we used Cosi2 [14] as haplotype simulation tool with parameters calibrated to empirical human data. We ran it with default settings for European population. Three separate experiments were conducted based on the total number of simulated haplotypes generated, viz. 1000 haplotypes, 5000 haplotypes and 10,000 haplotypes. Haplotypes in the collection had *l* = 1760 markers. Each simulated data set has a recombination rate sampled from a distribution matching the decode map[10] with recombination clustered into hotspots. The simulation package can be obtained from http://www.broad.mit.edu/sfs/cosi.

Simulated data were devoid of SNP ids. Hence public reference haplotype datasets were not included in these experiments. We used leave-one-out cross-validation (LOOCV) where known set *A_n_* is created by including all haplotypes except one pair and we infer constituent haplotypes from the combined genotype of the remaining pair. We ran these experiments with GPhase and with SHAPEIT. Default recombination settings were used for SHAPEIT. Mean switch accuracy results for GPhase and SHAPEIT for these cross validation experiments are given in table 1. GPhase phasing exhibited comparable or improved quality for all three values of *n*.

**Table 1.** Average percentage switch accuracy for cross validation experiments on simulated datasets with different sizes *n*.

| | $n = 1000$ | $n = 5000$ | $n = 10,000$ |
| --- | --- | --- | --- |
| GPhase | 97.51 | 97.5 | 98.73 |
| SHAPEIT | 96.1 | 97.2 | 98.16 |

We note that genotype samples considered in this experiment consists of about 1700 markers which is more than the typical number of markers seen at gene level. GPhase is able to obtain good phasing accuracy here as well even without explicitly modeling recombination. We believe that this is due to sufficiently large collection of known haplotypes *A_n_* in these experiments as larger collections would contain several haplotype instances resulting from different recombination possibilities. This perhaps alleviates the need for explicitly factoring recombination into the model.

We also performed leave-one-out cross-validation on 1000 genome data using GPhase. The data consisted of 5008 haplotypes each with 4859 markers. As in earlier experiment, all haplotypes except one pair were provided as *A_n_* in each cross-validation experiment. GPhase achieved mean switch accuracy of 97. 82 percentage in this case.

As discussed earlier, GPhase has linear run time and uses linear space. The tool implementation was done in Python. The tool does fast phasing and all inputs were phased in a matter of seconds.

## IV. Conclusion

We considered the problem of phasing an individual genotype sample given a collection of known haplotypes. We give a greedy approximation algorithm GPhase that uses the coalescent based mutation model of Stephens and Donnelly [16], which is also used in PHASE (v1.1) [18], for inferring a consistent haplotype pair. Though computing optimal solution under this model is computationally expensive, our algorithm computes an approximate solution efficiently in linear time and space. Experiments on real datasets revealed improved gene level phasing accuracy for GPhase tool compared to widely used tools such as SHAPEIT, Beagle, MaCH and Impute2. On simulated data, GPhase tool was able phase samples containing more than 1700 markers with high accuracy. GPhase tool can be used for gene level phasing of individual samples using publicly available haplotype datasets such as 1000 genome data. This has applications in studies on recessive Mendelian disorders in the absence of parent data. It would be interesting to provide theoretical bounds on the quality of the solution computed by GPhase algorithm, perhaps under certain assumptions on the known haplotype collection, that explains empirical findings. It would also be worthwhile to conduct chromosome level phasing experiments using GPhase with large collections of known haplotypes to evaluate its ability to deal with recombinations.

